# RAS G-domains fine-tune the sorting of phosphatidylserine acyl chains in the plasma membrane

**DOI:** 10.1101/2023.06.20.545612

**Authors:** Neha Arora, Hong Liang, Yong Zhou

## Abstract

Mutant RAS are major contributors to cancer and signal from nanoclusters on the plasma membrane (PM), via isoform-specific membrane anchors. However, the same RAS isoform bound to different guanine nucleotides are segregated on the PM. Paradoxically, various segregated RAS nanoclusters all enrich a type of anionic phospholipid, phosphatidylserine (PS). These findings suggest intricate participation of RAS G-domains in their PM distribution, which have not been explored. We now show that wild-types, oncogenic G12V mutants and membrane anchors of isoforms HRAS, KRAS4A and KRAS4B sort distinct PS species. Mechanistically, shifting orientation states of KRAS4B G-domain exposes residues, such as Arg 73, Arg 102 and Arg 135, to the PM, and contributes to PS acyl chain sorting. Oncogenic mutations may shift orientation states of G-domains. We show that G12V, G12D, G12C, G13D and Q61H mutants of KRAS4B sort distinct PS species. Thus, RAS G-domains fine-tune their lateral distribution on the PM.

## Introduction

RAS small GTPases, including isoforms HRAS, NRAS, splice variants KRAS4A and KRAS4B, toggle between the active GTP-bound and the inactive GDP-bound states, and participate in cell growth, division, survival, proliferation, and migration ^1–3^. Mutant RAS at key residues, such as glycine 12 (Gly 12), Gly 13 and glutamine 61 (Gln 61), remain constitutively active and are main drivers of cancer ^1–3^. Signaling of wild-type and oncogenic mutants of RAS proteins is mostly compartmentalized to spatially distinct nano-domains, termed as nanoclusters, on the plasma membrane (PM) ^3–6^. Their isoform-specific C-terminal membrane-anchoring domains have been attributed to mainly facilitate association with lipids and the formation of nanoclusters on the PM ^3–8^. Interestingly, it has long been observed that, with the identical membrane anchor, the same RAS isoform in active and inactive states are spatially segregated on the PM ^9–11^. These findings strongly suggest that RAS G-domains, mostly suspended in the cytoplasm, contribute to lipid sorting. If their G-domains play roles in lipid sorting, will various oncogenic mutations of RAS G-domains differentially alter lipid sorting? These aspects of RAS function have not been explored before. Further, despite being spatially segregated, all RAS nanoclusters tested enrich the same type of anionic phospholipid, phosphatidylserine (PS) ^11–15^. This finding implies that different parallel pools of PS co-exist among various RAS nanoclusters on the PM. Indeed, we previously showed that the nanoclusters of oncogenic mutant KRAS4B^G12V^ selectively enriched the mixed-chain PS species with one saturated and one unsaturated acyl chains, but not other PS species with dual saturated or dual unsaturated acyl chains ^12,13,15^. We further showed the efficient recruitment of effector CRAF by the constitutively active KRAS4B^G12V^ only occurred in the presence of the mixed-chain PS species, but not other symmetric PS species tested ^12^. Intact caveolae also selectively enrich the symmetric PS species, but not the mixed-chain PS ^16^. Parallel cholesterol-dependent and cholesterol-independent PS pools have also been shown to co-exist in the PM ^13,17^. Yet, do RAS G-domains, which do not have direct access to the bilayer core, participate in the sorting of lipid acyl chains? Here, our current study aims to systematically examine whether and how RAS G-domains selective sort PS species in cells.

Via super-resolution electron microscopy (EM)-spatial analysis, we here show that the full-length wild-types, oncogenic G12V mutants and the truncated minimal membrane-anchoring domains of HRAS, KRAS4A and KRAS4B sort distinct PS species on the PM. These data strongly suggest that the G-domains of RAS isoforms contribute to the selectivity of PS acyl chains. Mechanistically, the G-domain of KRAS4B has been predicted to sample distinct orientation states on the PM. As a result, KRAS4B G-domain possesses distinct membrane-interacting interfaces during these orientation states. To explore mechanisms, we mutated key residues in the membrane-interacting interfaces of the G-domain of KRAS4B, including Arginine 73 (Arg 73), Arg 102 and Arg 135. We show that mutations of G-domain residues differentially alter the select sorting of PS species of the wild-type and G12V mutant of KRAS4B. Oncogenic mutations of KRAS4B also shift the reorientation states of its G-domain, we further compared the PS sorting of oncogenic mutants G12V, G12C, G12D, G13D and Q61H of KRAS4B. We now show that these oncogenic mutants of KRAS4B enrich different profiles of PS species. Thus, G-domains of RAS isoforms contribute to the select sorting of PS acyl chains and fine-tune the lateral segregation of RAS isoforms on the PM. Our data also suggest that, despite being similarly constitutively active, KRAS4B oncogenic mutants may occupy non-overlapping spaces on the PM, which may contribute to their mutant-specific pathological activities.

## Results

### RAS G-domains contribute to the select sorting of PS species

To examine the potential roles of RAS G-domains in facilitation of lipid sorting, we first compared how the full-length wild-types, oncogenic G12V mutants and the minimal membrane anchors of isoforms HRAS, KRAS4A and KRAS4B sorted different PS species. Typical mammalian cells contain 30-40 PS species. In the current study, we compared 3 PS species: the fully saturated di18:0 PS (DSPS), the mono-unsaturated di18:0 PS (DOPS) or the mixed-chain 18:0/18:1 PS (SOPS), as these species represent major classes of lipid acyl chains. To modulate PS contents in cells, PS auxotroph (PSA3) cells were grown in medium containing 10% dialyzed fetal bovine serum (DFBS) for 72 hours to deplete endogenous PS ^11–18^. To restore the normal endogenous PS levels, PSA3 cells were supplemented with 10 μM ethanolamine (Etn) for 72 hours ^11–18^. To compare effects of individual PS species, we acutely added back synthetic PS species to the PSA3 cells depleted of endogenous PS via 1 hour incubation with synthetic PS species ^12,13,15^. We used electron microscopy (EM)-bivariate co-clustering analysis to quantify the co-localization between GFP-LactC2 (a PS-specific binding domain) and an RFP-tagged RAS construct on intact PM sheets of PSA3 cells. The EM-bivariate co-clustering analysis has been extensively used to quantify co-localization of PM constituents ^11–16^. Briefly, the apical PM sheets of PSA3 cells were attached to copper EM grids. Following fixation, the PM sheets were immunolabeled with 6 nm gold nanoparticles conjugated to anti-GFP antibody and 2 nm gold nanoparticles coupled to anti-RFP antibody, respectively. The gold distribution on the intact PM sheets was imaged via transmission EM (TEM) at 100,000X magnification. Co-clustering between 6 nm and 2 nm gold particles within a 1 μm^2^ PM area was calculated using the Ripley’s bivariate K-function analysis. The extent of co-clustering, *L_biv_*(*r*) – *r*, was plotted as a function of distance *r* in nanometers. The area-under-the-curve for the bivariate K-function curves between the *r* values of 10 nm and 110 nm was termed as L-function bivariate integrated (LBI). The LBI value of 100 is the 95% confidence interval (95% CI), the values above which indicate statistically significant co-clustering between the two gold populations. The larger LBI values correspond to more extensive co-clustering and higher enrichment.

Fig.1A shows that the LBI value between GPF-LactC2 and RFP-KRAS4B^G12V^ was ~178 in PSA3 cells supplemented with Etn, illustrating extensive co-clustering between the oncogenic mutant KRAS4B^G12V^ and the endogenous PS in the PM of PSA3 cells containing the normal endogenous PS level. Depletion of endogenous PS (DFBS) significantly decreased the LBI value below the 95% C.I, suggesting efficient spatial segregation between KRAS4B^G12V^ and the endogenous PS upon PS depletion. In the PS-depleted PSA3 cells, acute addback of SOPS, but not other PS species tested, effectively restored the co-clustering between GFP-LactC2 and RFP-KRAS4B^G12V^. This set of data was entirely consistent with our previous findings that the PM nanoclusters of mutant KRAS4B^G12V^ selectively enriched the mixed-chain PS species ^12,13,15^. Similar to RFP-KRAS4B^G12V^, the wild-type RFP-KRAS4B and the minimal anchor RFP-tK(4B) also co-clustered extensively with the endogenous PS (Etn, Fig.1B and C). Depletion of endogenous PS (DFBS) segregated GFP-LactC2 from RFP-KRAS4B or RFP-tK(4B). Interestingly, the wild-type RFP-KRAS4B co-localized with DOPS and SOPS, but not DSPS (Fig.1B). The minimal anchor RFP-tK(4B) co-clustered with all 3 PS species tested, while associating most extensively with SOPS (Fig.1C). Thus, while all 3 KRAS4B constructs preferentially associate with the mixed-chain PS, the mutant KRAS4B^G12V^ possesses more selectivity.

**Figure 1.**
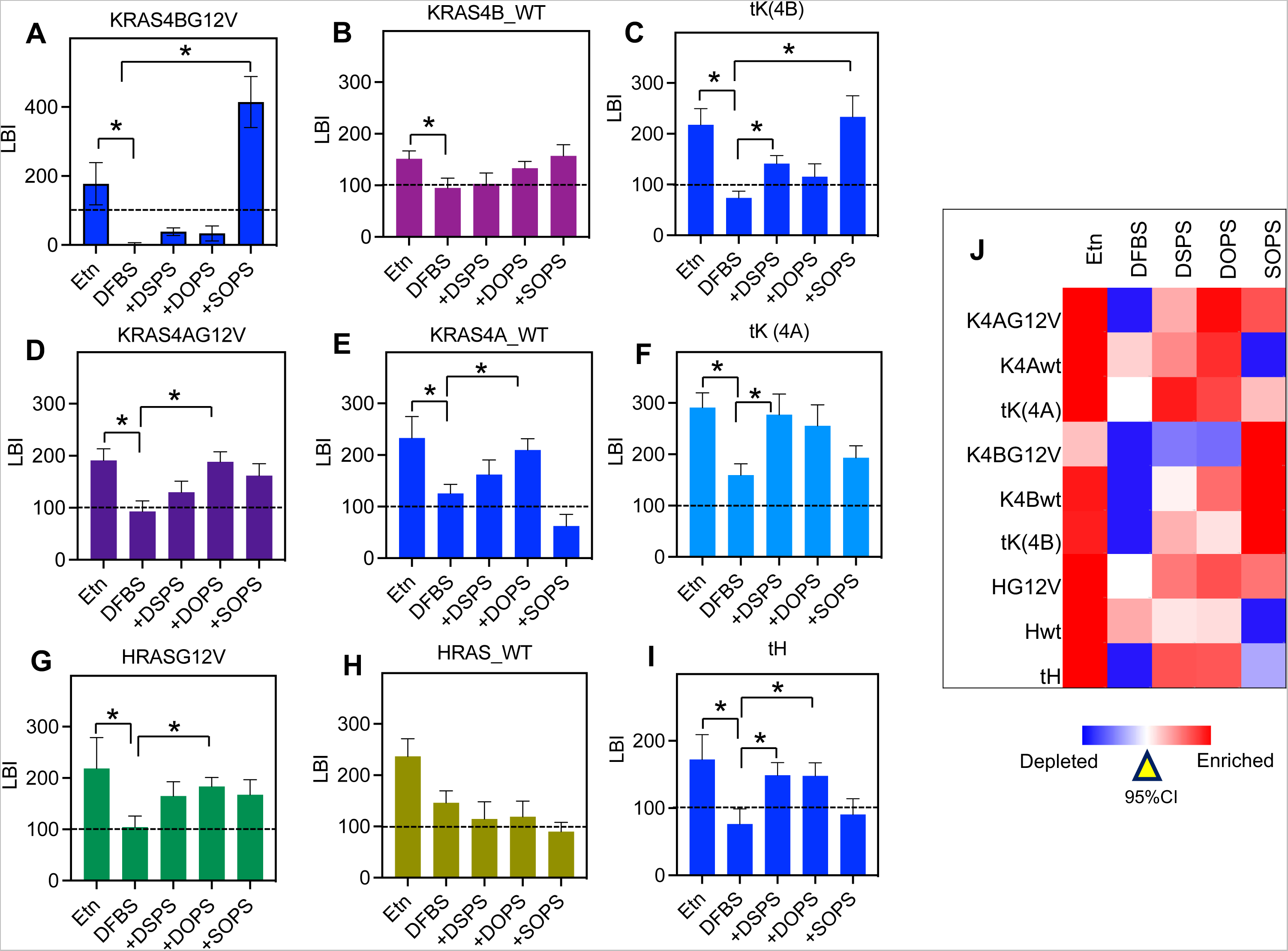
Wild-types, G12V mutants and minimal anchors of RAS isoforms sort distinct PS species. Endogenous PS in the PS auxotroph PSA3 cells was depleted via growing the cells in 10% dialyzed FBS (DFBS) for 72 hours. The normal endogenous PS was restored via supplementation of 10 μM ethanolamine (Etn) for 72 hours. To compare individual PS species, exogenous PS species was acutely added back to the PS-depleted PSA3 cells. Specifically, the PS-depleted PSA3 cells grown in DFBS were incubated with the medium containing 10 μM di18:0 PS (DSPS), di18:1 PS (DOPS) or 18:0/18:1 PS (SOPS) for 1 hour before harvesting. PSA3 cells were ectopically co-expressing GFP-LactC2 (a PS probe) and an RFP-tagged RAS construct: RFP-tagged KRAS4B constructs (A-C), RFP-tagged KRAS4A (D-F) and RFP-tagged HRAS constructs (G-I). Intact apical PM sheets of PSA3 cells were attached to copper EM grids. GFP and RFP anchored to the PM inner leaflet were immunolabeled with 6nm gold nanoparticles conjugated to anti-GFP antibody and 2nm gold coupled to anti-RFP antibody, respectively. EM-bivariate co-clustering analysis quantified the co-clustering between the 6nm and 2nm gold populations, with LBI values indicating the extent of co-clustering or enrichment. The LBI value of 100 is the 95% confidence interval (95% CI, the dashed lines), the values above which indicate statistically significant co-clustering. For each condition, at least 15 apical PM sheets from individual cells were imaged, analyzed, and pooled together. Data are shown as mean ± SEM. Statistical significance was evaluated using bootstrap tests, with * indicating p < 0.05.

The splice variants KRAS4A and KRAS4B differ significantly in their C-terminal membrane-anchoring domains, with KRAS4A possessing an additional palmitoyl chain without the polybasic domain. We previously showed that an oncogenic mutant KRAS4A^G12V^ also co-clustered with the endogenous PS ^19^. Consistently, Fig.1D-F show that KRAS4A^G12V^, wild-type KRAS4A, and the minimal anchor tK(4A) all enriched endogenous PS extensively (Etn). PS depletion (DFBS) effectively separated the PS probe from KRAS4A constructs (Fig.1D-F). For the PS acyl chain specificity, RFP-KRAS4A^G12V^ co-clustered with all 3 PS species tested. The wild-type RFP-KRAS4A excluded SOPS, while RFP-tK(4A) co-clustered with all 3 PS species with the highest extent of co-clustering with DSPS. Taken together, KRAS4A constructs sort PS acyl chains in distinct manners.

We next compared the PS acyl chain sorting of isoform HRAS. Consistent with previous findings, Fig.1G-I show that the oncogenic mutant HRAS^G12V^, the full-length HRAS and the minimal anchor tH, all extensively enriched with the endogenous PS (Etn) and dissociated from PS upon PS depletion (DFBS) ^11,14^. When comparing individual PS species, RFP-HRAS^G12V^ co-clustered equivalently with all 3 PS species tested (Fig.1G). Interestingly, the co-clustering LBI parameter between GFP-LactC2 and the wild-type RFP-HRAS was near 95% CI for all 3 PS species tested. This data suggests that the wild-type HRAS does not significantly co-localize with all 3 PS species tested. The minimal anchor RFP-tH more preferentially associated with DSPS and DOPS, but not SOPS, consistent with previous molecular dynamic simulations that tH preferentially localizes to the boundaries between the liquid-ordered domains (*L_o_* domains enriching saturated lipids) and the liquid-disordered domains (*L_d_*domains containing unsaturated lipids) ^20^. Taken together and summarized in a heatmap of LBI values (Fig.1J), RAS isoforms selectively sort distinct profiles of PS species in isoform- and guanine nucleotide-specific manners, strongly suggesting that G-domains participate in the selectivity of lipid acyl chains.

### KRAS4B G-domain residues contribute to the select sorting of PS acyl chains

It is puzzling how these largely cytosolic G-domains of small GTPases might recognize lipid acyl chains hidden within the membrane bilayer core. To examine this question mechanistically, we focused on KRAS4B since the structures and conformational orientation of the G-domain of KRAS4B on membranes have been extensively studied. Previous all-atom molecular dynamic (AA-MD) simulations proposed that the G-domain of KRAS4B switches between two distinct orientation states, OS1 and OS2, as well as an intermediate OS0 state ^21,22^. The mutant KRAS4B^G12V^ has been predicted to disfavor the OS2 state ^21^. Specific basic residues of the G-domain of KRAS4B, such as Arg 135 in OS1 and Arg 73 and Arg 102 in OS2, form close contacts with lipid bilayers at high probability ^21–23^. In consequence, the C-terminal membrane anchor of KRAS4B alters conformations in coordination with the reorientation of its G-domain. KRAS4B membrane anchor reorients to face the lobe 2 (residues 87-166) in OS1 and turns to face the lobe 1 (residues 1-86) in OS2 ^21,22^. Thus, the intercalation of KRAS4B membrane anchor into membranes may coordinate with the reorientation of its G-domain, thus altering the sorting of PS acyl chains. To examine how G-domain reorientation participate in PS acyl chain sorting, we mutated Arg 73, Arg 102 or Arg 135 of the wild-type KRAS4B and the oncogenic mutant KRAS4B^G12V^ and generated a cohort of OS mutants: RFP-KRAS4B^R73Q^, RFP-KRAS4B^R102Q^, RFP-KRAS4B^R135Q^, RFP-KRAS4B^G12V.R73Q^, RFP-KRAS4B^G12V.R102Q^ and RFP-KRAS4B^G12V.R135Q^. We performed EM-bivariate co-clustering analysis following acute addback of various PS species to PSA3 cells depleted of endogenous PS. Unlike the original G-domain of RFP-KRAS4B^G12V^, RFP-KRAS4B^G12V.R73Q^ no longer co-clustered with the endogenous PS (Etn) (Fig.2A and B). Interestingly, RFP-KRAS4B^G12V.R73Q^ co-clustered extensively with DSPS (Fig. 2B). RFP-KRAS4B^G12V.R102Q^ behaved similarly to the original RFP-KRAS4B^G12V^ and preferentially sorted SOPS (Fig.2C). RFP-KRAS4B^G12V.R135Q^ preferred DOPS and then DSPS, but not SOPS (Fig.2D).

**Figure 2.**
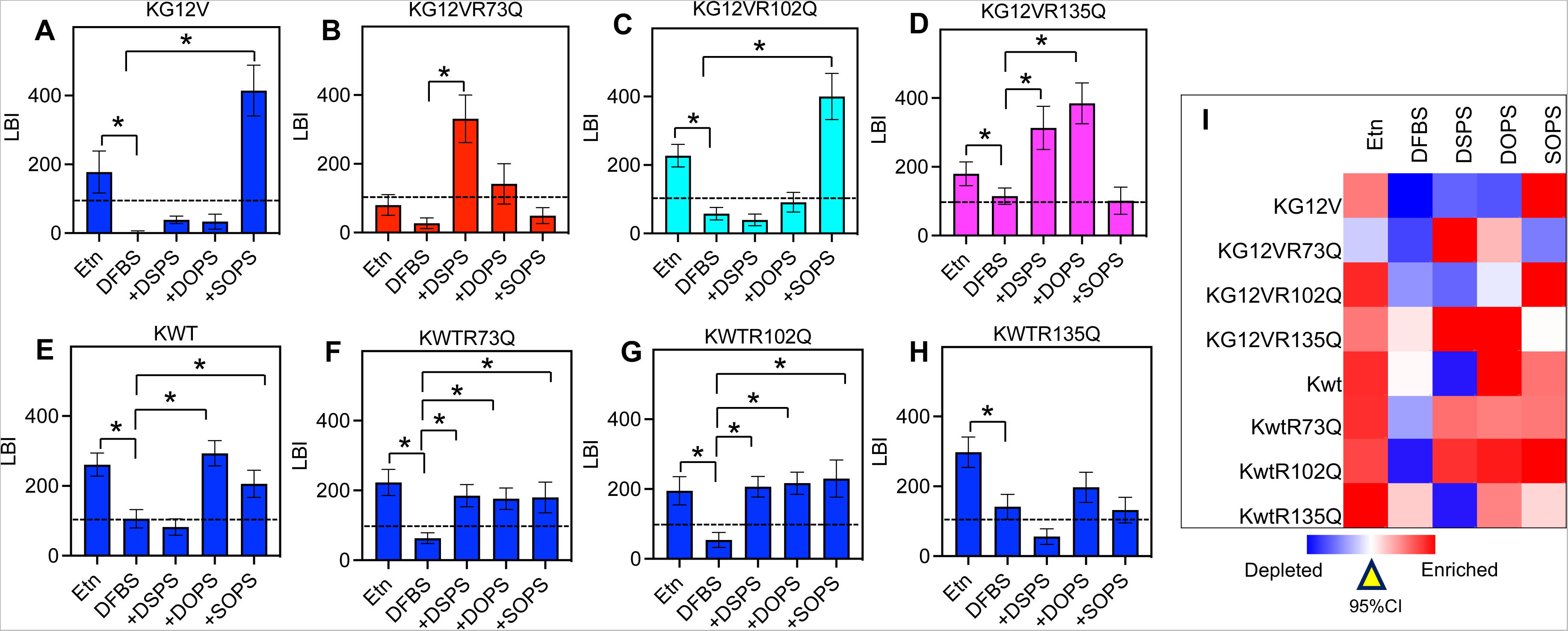
Key KRAS4B G-domain residues at the membrane interface contribute to the select sorting of PS acyl chains. Key membrane-associating residues, such as Arg 73, Arg 102 and Arg 135, were mutated in the oncogenic KRAS4B^G12^V (A-D) and the wild-type KRAS4B (E-H) to generate a cohort of orientation state (OS) mutants. PSA3 cells co-expressing GFP-LactC2 and an RFP-tagged OS mutant were supplemented with Etn, DFBS and/or individual exogenous PS species (DSPS = di18:0 PS, DOSP = di18:1 PS, SOPS = 18:0/18:1 PS) as described in Fig.1. EM-bivariate analysis quantified the co-clustering or enrichment of endogenous or exogenous PS species within the nanoclusters of OS mutants of KRAS4B. For each condition, at least 15 PM sheets from individual cells were imaged, analyzed and pooled. LBI values are shown as mean ± SEM. Bootstrap tests evaluated the statistical significance between data sets.

The OS mutants of the wild-type RFP-KRAS4B displayed different patterns of PS acyl chain selectivity. RFP-KRAS4B^R73Q^ and RFP-KRAS4B^R102Q^ associated with all 3 PS species tested, suggesting that mutating Arg 73 and Arg 102 in OS2 of the wild-type KRAS4B lost selectivity for acyl chains of PS, while still maintaining high affinities for the PS headgroup (Fig.2E-G). Acute addback of any of the three PS species did not effectively restore the co-clustering between GFP-LactC2 and RFP-KRAS4B^R135Q^ when compared with the PS-depleted condition (DFBS) (Fig.2H). This suggests that KRAS4B^R135Q^ does not associate with any of the PS species tested. The significant differences between the OS mutants of KRAS4B^G12V^ and those of the wild-type KRAS4B strongly suggest that reorientation of KRAS4B G-domain contributes to the selectivity of PS acyl chains (summarized in a heatmap of LBI values in Fig.2I).

### KRAS4B oncogenic mutants selectively sort distinct PS species

KRAS4B oncogenic mutations in residues Gly 12, Gly 13 and Gln 61, such as G12D, G12V, G12C, G13D and Q61H, result in prolonged GTP-binding and are frequently found in cancer ^1–3^. Although these residues are not proximal to membranes, their mutations have been predicted to shift the reorientation equilibrium between OS1 and OS2 ^24–26^. Thus, it is possible that these oncogenic mutants may sort distinct PS species. To test this, we performed similar EM-bivariate co-clustering analysis and compared co-clustering between GFP-LactC2 and an RFP-tagged KRAS4B oncogenic mutant in PSA3 cells. Shown in Fig.3, KRAS4B^G12D^ and KRAS4B^Q61H^ behaved similarly with KRAS4B^G12V^ (Figs1 and 2), and preferentially enriched with SOPS but not other PS species tested. Interestingly, KRAS4B^G12C^ associated with all 3 PS species tested, while KRAS4B^G13D^ more preferentially associated with the symmetric DSPS and DOPS, but not SOPS. Taken together, although oncogenic mutants of KRAS4B are all constitutively active, they possess distinct capabilities to sort different PS species (summarized in a heatmap of LBI values in Fig.3E).

**Figure 3.**
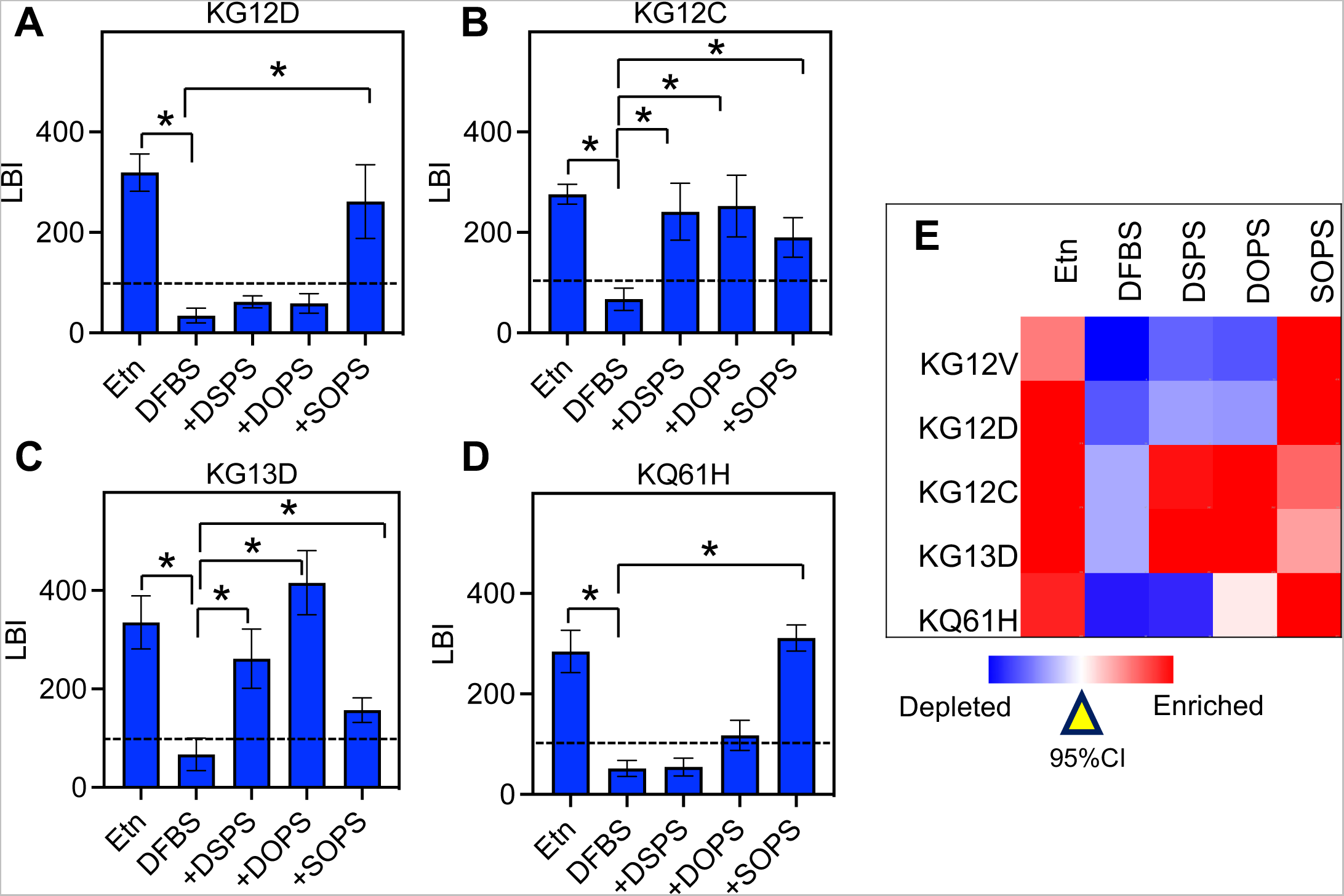
Oncogenic mutants of KRAS4B selectively enrich distinct profiles of PS species. PS contents in the PSA3 cells co-expressing GFP-LactC2 and an RFP-tagged oncogenic mutant of KRAS4B, such as KRAS4BG12D (A), KRAS4BG12C (B), KRAS4BG13D (C) and KRAS4BQ61H (D) were modulated as described above. Etn = normal endogenous PS levels, DFBS = endogenous PS depletion, DSPS = di18:0 PS, DOSP = di18:1 PS, SOPS = 18:0/18:1 PS. Exogenous PS was acutely added back to the PSA3 cells depleted of endogenous PS. EM-bivariate co-clustering analysis calculated the co-clustering LBI values and quantified the enrichment of PS species within the nanoclusters of KRAS4B oncogenic mutants. The LBI value of 100 is the 95%CI. For each condition, at least 15 apical PM sheets were imaged, analyzed and pooled. LBI values are shown as mean ± SEM. Bootstrap tests evaluated the statistical significance between data sets.

## Discussion

Although the isoform-specific C-terminal membrane anchors of RAS proteins have been primarily focused for their membrane interactions, it has long been observed that the same RAS isoform bound to different guanine nucleotides occupy non-overlapping spaces on the PM ^9,10^, strongly suggesting that RAS G-domains play roles in their lateral distribution on the PM. Indeed, atomistic MD simulations and fluorescence lifetime imaging microscopy - fluorescence resonance energy transfer (FLIM-FRET) demonstrated that the G-domain of HRAS undergoes significant reorientation upon GTP-GDP exchange, and in turn alters the insertion depth of its dual palmitoyl chains and farnesyl chain in the membranes ^27,28^. This consequentially results in the preferential partitioning of the GTP-bound HRAS to the cholesterol-independent domains and the GDP-bound HRAS to the cholesterol-enriched domains ^27–29^. However, the G-domain of KRAS4B is more dynamic and does not undergo the profound structural reorientation as the G-domain of HRAS. Despite that, all-atom MD simulations and single molecule-fluorescence resonance energy transfer (smFRET) proposed that KRAS4B G-domain adopts distinct conformational orientations and dynamically interact with polar components at the membrane surface ^21–23,30^. In particular, the G-domain of KRAS4B can be described as having 2 main lobes: lobe 1 contains residues 1-86 and lobe 2 contains residues 87-166. The α3-5 helices (on lobe 2) in the orientation state 1 (OS1) and the β1-3 loops (on lobe 1) in the OS2 contact the membranes, with an additional intermediate OS0 state (Fig.4A). A cohort of charged residues has been proposed to interact with PS headgroup, including Arg 97, Glu 98, Lys 101, Glu 107, Asp 108, Lys 128, Asp 132 and Arg 135 in OS1, and Arg 73 and Arg 102 in OS2 ^21–23,30^. KRAS4B G-domain residues, such as Tyr 64, Ser 65, Arg 68, Tyr 71 and Arg 73, also extensively associate with the zwitterionic phosphatidylcholine (PC) lipids ^22,23^. As illustrated in Fig.4A, the membrane anchor of KRAS4B adopts distinct conformations in different OS states. Thus, changing equilibrium among various OS states may in coordination alter interactions between KRAS4B anchor and lipids. Our current study shows that mutating residues at the membrane-interacting interface, such as Arg 73, Arg 102 and Arg 135, significantly alters the sorting of PS species by the wild-type KRAS4B and oncogenic mutant KRAS4B^G12V^. Especially, it has been proposed that oncogenic mutant KRAS4B^G12V^ disfavors OS2 ^21^. This is consistent with our finding that mutating Arg 135, which associates with PS extensively in OS1, most profoundly altered the sorting of PS species (Fig.2D). On the other hand, mutating Arg 102 of KRAS4B^G12V^ did not impact the PS acyl chain sorting (Fig.2C) since Arg 102 associates with PS in OS2. Thus, RAS G-domains and their C-terminal membrane anchors may coordinate to synergistically participate in the intricate sensing of local environments in the PM.

**Figure 4.**
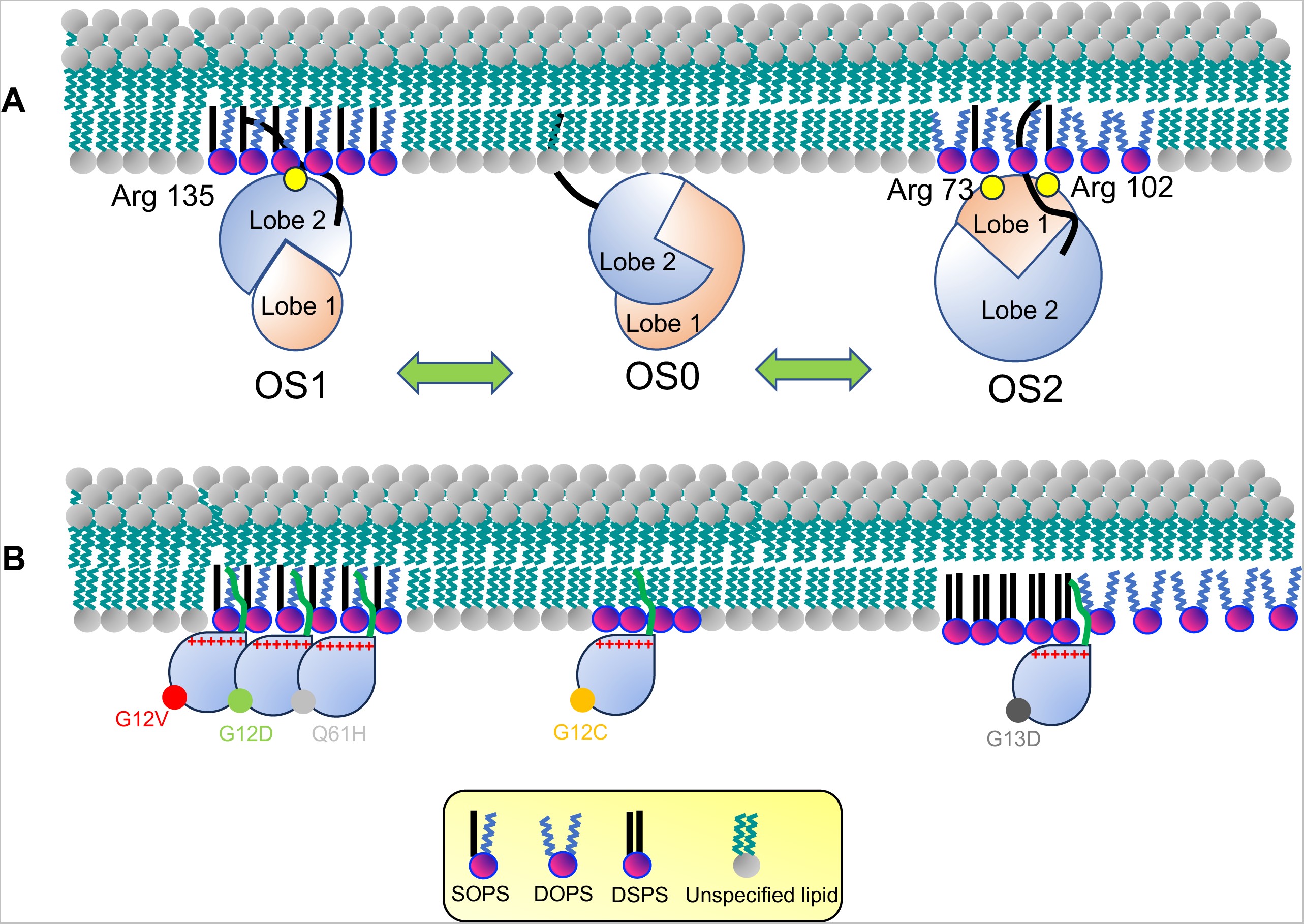
KRAS4B G-domain switches among orientation states on the membrane and expose distinct basic residues to lipids. (A) A schematic illustration describes dynamic equilibrium among 3 orientation states for the G-domain of KRAS4B, OS1 ↔ OS0 and OS0 ↔ OS2. OS1 and OS2 states do not switch between each other and they all go through OS0 as an intermediate state. Distinct membrane-associating surfaces expose to the membranes at different OS states and present distinct sets of basic residues to associate with lipids, such as Arg 135 in OS1, and Arg 73 and Arg 102 in OS2. Oncogenic mutations, although not proximal to membranes, alter the equilibrium among OS states and in turn modulate the frequencies of exposure of these basic residues to lipids. (B) Various oncogenic mutants of KRAS4B laterally distribute to distinct regions on the PM.

Additional mechanisms can also contribute to how RAS G-domains detect lipid acyl chain structures. RAS G-domains specifically interact with complex constituents, such as RAF/PI3K effectors, actin, galectins, and chaperones such as phosphodiesterase (PDEδ) and/or a G protein-coupled receptor GPR31 ^10,31–37^. Indeed, the formation of signalosomes composed of KRAS4B, effectors BRAF CRAF, downstream kinase MEK, galectin-3 (Gal-3) and 14-3-3 exposes G-domain α4 and α5 helices to the membranes ^38^. These signaling platforms may also contribute to how RAS G-domains participate in the sorting of lipid acyl chains. Taken together, the complex nanoclusters and signaling platforms of RAS may integrate a series of weak protein/protein and protein/membrane interactions and allow synergistic regulation of the select sorting of lipid acyl chains.

Oncogenic mutations of RAS at Gly 12, Gly 13 and Gln 61 play important roles in cancer. Although these mutations all deactivate the GTPase activities of RAS and allow RAS to remain constitutively active for prolonged periods, their oncogenic activities differ significantly. While the G12 residue of KRAS4B is most mutated in cancer (81%), the Q61 residue of HRAS is most mutated (38%) ^2^. The G12D, G12V and G12C mutations are found in 46%, 31% and 3% of KRAS4B oncogenic mutations in pancreatic ductal adenocarcinoma, respectively ^39^. On the other hand, the G12C mutation is found in 37% of KRAS4B mutations in non-small cell lung cancer ^39^. KRAS4B^G12V^ is more efficient than KRAS4B^G12D^ in the activation of the mitogen-activated protein kinases (MAPKs) cascade ^40^. These observations strongly suggest that these constitutively active mutants possess distinct biological activities. Here, we show that KRAS4B^G12V^, KRAS4B^G12D^, KRAS4B^Q61H^ share similar affinities for the mixed-chain PS species, while KRAS4B^G12C^ may have lost selectivity for PS acyl chains but still maintaining high affinities for PS headgroup (Fig.4B). Interestingly, KRAS4B^G13D^ prefers the saturated DSPS and the unsaturated DOPS (Fig.3C). The saturated lipids typically favor the cholesterol-enriched rafts, while the unsaturated lipids favor the cholesterol-poor non-raft domains. Our data suggest that KRAS4B^G13D^ may distribute to the boundary regions between the rafts and non-rafts. Indeed, HRAS membrane anchor tH has been predicted to localize to the raft/non-raft boundaries ^20^ and favorably associate with both the saturated PS and unsaturated PS (Fig.1I). Although we focused on the PS species in the current study, it is possible that some KRAS4B mutants, such as KRAS4B^G12C^ and KRAS4B^G13D^, may enrich raft lipids, such as PIP_2_ and cholesterol. Thus, in tumor cells expressing multiple oncogenic mutants of KRAS4B, some mutants, such as G12V, G12D and Q61H, may co-localize together and signaling from the same nanoclusters on the PM. Other mutants, such as KRAS4B^G12C^ and KRAS4B^G13D^, may segregate to distinct nano-domains.

## Conclusion

Mutant RAS proteins primarily signal from their spatially distinct nanoclusters on the PM, largely mediated via associations of their membrane anchors and PM lipids. Here, we show exciting evidence that RAS G-domains, mostly suspended in the cytoplasm, also participate in the intricate sensing of lipid acyl chains. These selective associations allow the same RAS isoforms bound to different guanine nucleotides to spatially segregate on the PM. Similarly, the G-domain-mediated sorting of lipid acyl chains allows various oncogenic mutants of RAS to either co-localize together or segregate to independent nano-domains on the PM. Since effector recruitment and the formation of signaling platforms require distinct lipid environment in the PM, these mechanisms revealed in the current study may contribute to the mutant-specific pathological activities of these mutants.

## Acknowledgement

This work was supported in part by the National Institutes of Health (NIH) R01GM138668.

## Materials and Methods

### Electron microscopy (EM)-spatial analysis

#### EM-univariate nanoclustering

The K-function univariate analysis quantifies the extent of nanoclustering of 4.5 nm gold nanoparticles conjugated to primary antibodies against a targeted protein on intact plasma membrane (PM) sheets ^12,14^. To probe the nanoclustering of a GFP-tagged RAS, the apical PM of PS auxotroph PSA3 cells ectopically expressing GFP-tagged RAS was attached to copper EM grids. Following fixation with 4% paraformaldehyde (PFA) and 0.1% gluaraldehyde, GFP on the PM sheets was immunolabeled with 4.5nm gold nanoparticles conjugated to anti-GFP antibody and embedded in methyl cellulose containing 0.3% uranyl acetate. Gold distribution on the intact PM sheets was imaged using TEM at 100,000x magnification. The coordinates of every gold particle were assigned via ImageJ. Nanoclustering of gold particles within a selected 1μm^2^ PM area was quantified using Ripley’s K-function under a null hypothesis that the point pattern in a selected area is distributed randomly:

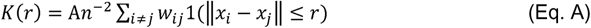

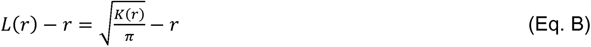

where *K(r*) indicates the univariate K-function for *n* gold nanoparticles in an intact PM area of *A*; *r* is the length scale between 1 and 240 nm with an increment of 1 nm; || · || is Euclidean distance; where the indicator function of 1(·) = 1 if ||*x_i_*−*x_j_*|| ≤ r and 1(·) = 0 if ||*x_i_*−*x_j_*|| > r. To achieve an unbiased edge correction, a parameter of *w*_ij_ ^−1^ is used to describe the proportion of the circumference of a circle that has the center at *x_i_* and radius ||*x_i_*−*x_j_*||. *K*(*r*) is then linearly transformed into *L*(*r*) – *r*, which is normalized against the 99% confidence interval (99% C.I.) estimated from Monte Carlo simulations. A *L*(*r*) − *r* value of 0 for all values of *r* indicates a complete random distribution of gold. A *L*(*r*) − *r* value above the 99% C.I. of 1 at the corresponding value of *r* indicates statistical clustering at certain length scale. At least 15 PM sheets were imaged, analyzed and pooled for each condition in the current study. Statistical significance was evaluated via comparing our calculated point patterns against 1000 bootstrap samples in bootstrap tests ^12,14^.

#### EM-Bivariate co-clustering analysis

The K-function bivariate co-clustering analysis quantifies the co-clustering between two differently sized gold nanoparticles tagging two different constituents on the intact PM sheets ^12,14^. Similar to the univariate nanoclustering protocol described above, intact apical PM sheets of PSA3 cells co-expressing GFP-LactC2 (probing PS lipids) and an RFP-tagged RAS construct were attached to EM grids and fixed with 4% PFA and 0.1% gluaraldehyde. The PM sheets were incubated with 6 nm gold nanoparticles linked to anti-GFP antibody, blocked with 0.2% bovine serum albumin (BSA) and 0.2% fish skin gelatin, then incubated with 2 nm gold conjugated to anti-RFP antibody. ImageJ was used to assign coordinates to the gold nanoparticle. A bivariate K-function analysis tested the null hypothesis that the two populations of gold particles spatially separate from each other. (Eqs. C–F):

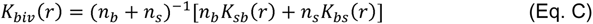

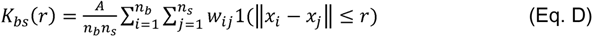

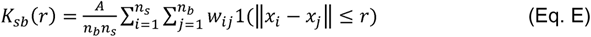

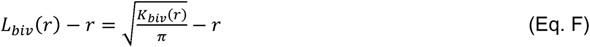

where *K_biv_*(*r*) denotes a bivariate estimator and contains two individual bivariate K-functions: *K_bs_*(*r*) quantifies how the big 6 nm gold particles (*b* = big gold) distribute around each 2 nm small gold particle (*s* = small gold); *K_sb_*(*r*) describes how small gold particles distribute around each big gold particle. The value of n_b_ indicates the number of 6 nm big gold and n_s_ indicates the number of 2nm small gold within a PM area of *A*. Other parameters denote the same definitions as defined in the univariate calculations in Eqs. A and B. *L_biv_*(*r*)−*r* is a linearly transformation of *K_biv_*(*r*), and is normalized against the 95% confidence interval (95% C.I.). An *L_biv_*(*r*)−*r* value of 0 indicates spatial segregation between the two populations of gold particles, whereas an *L_biv_*(*r*)−*r* value above the 95% C.I. of 1 at the corresponding distance of *r* indicates yields statistically significant co-localization at certain distance yields. Area-under-the-curve for each *L_biv_*(*r*)−*r* curves was calculated within a fixed range 10 < *r* < 110 nm, and was termed bivariate *L_biv_*(*r*)−*r* integrated (or LBI):

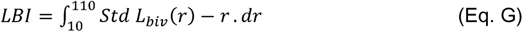

For each condition, > 15 apical PM sheets were imaged, analyzed and pooled, shown as mean of LBI values ± SEM. Statistical significance between conditions was evaluated via comparing against 1000 bootstrap samples as described ^12,14^.

