## Supplemental figure for "RAS G-domains fine-tune the sorting of phosphatidylserine acyl chains in the plasma membrane"

### Supplemental Figure 1

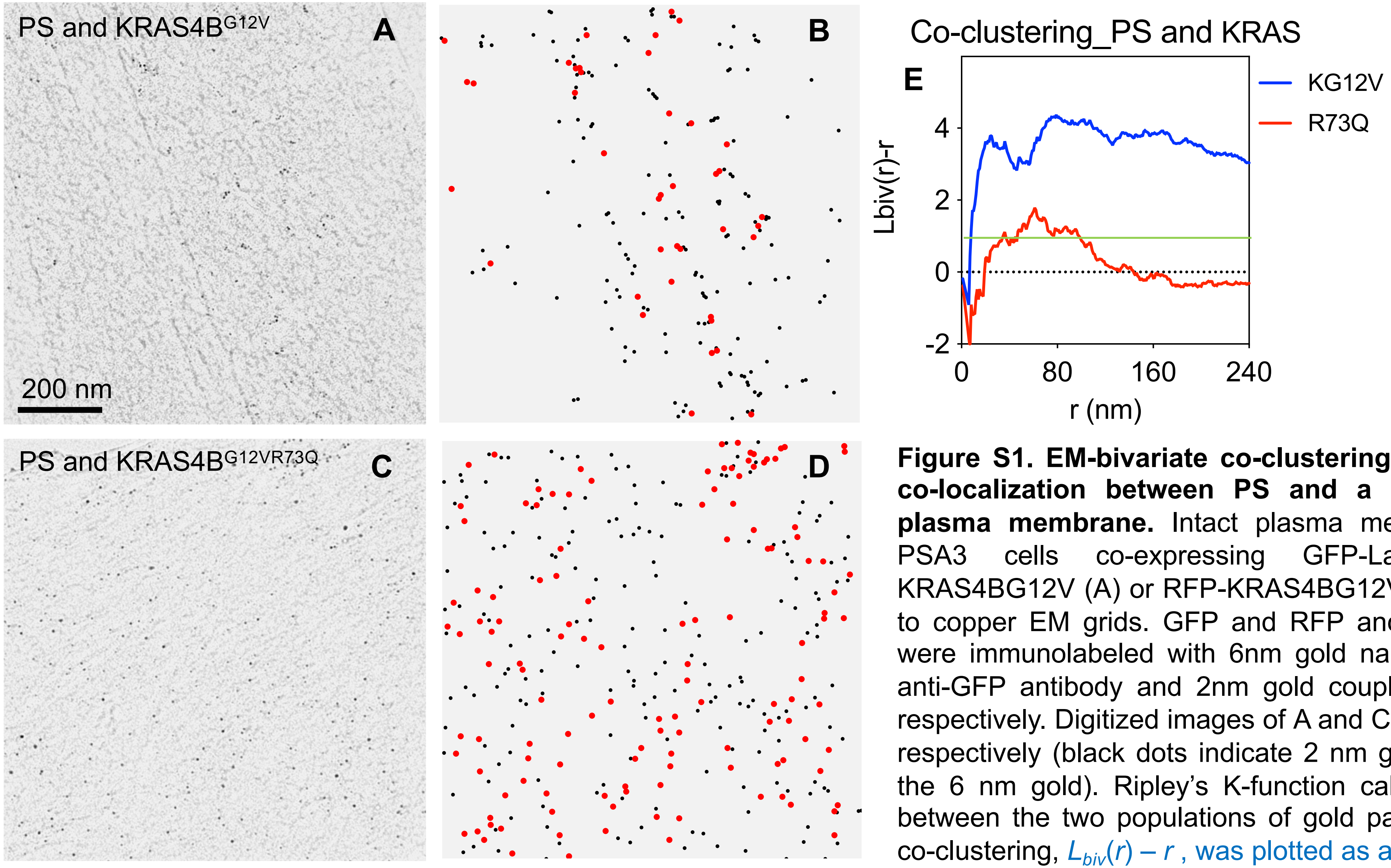


**Figure S1. EM-bivariate co-clustering analysis quantifies the co-localization between PS and a RAS construct on the plasma membrane.** Intact plasma membrane (PM) sheets of PSA3 cells co-expressing GFP-LactC2 and an RFP-KRAS4BG12V (A) or RFP-KRAS4BG12VR73Q (C) were attached to copper EM grids. GFP and RFP anchored to the PM sheets were immunolabeled with 6nm gold nanoparticles conjugated to anti-GFP antibody and 2nm gold coupled to anti-RFP antibody, respectively. Digitized images of A and C are shown in (B) and (D), respectively (black dots indicate 2 nm gold and red dots indicate the 6 nm gold). Ripley's K-function calculated the co-clustering between the two populations of gold particles. (E) The extent of co-clustering, $L_{biv}(r) - r$, was plotted as a function of length scale, $r$ in nanometer (nm). The $L_{biv}(r) - r$ curves for A and C are shown. The green line indicates the 95% confidence interval.
